# RoPE: a robust profile likelihood method for differential gene expression analysis

**DOI:** 10.1101/2022.06.02.494449

**Authors:** Lehang Zhong, Lisa J. Strug

## Abstract

Variation in RNA-Seq data creates modeling challenges for differential gene expression (DE) analysis. Statistical approaches address conventional small sample sizes and implement empirical Bayes or non-parametric tests, but frequently produce different conclusions. Increasing sample sizes enable proposal of alternative DE paradigms. Here we develop RoPE, which uses a data-driven adjustment for variation and a robust profile likelihood ratio DE test. Simulation studies show RoPE can have improved performance over existing tools as sample size increases and has the most reliable control of error rates. Application of RoPE demonstrates that an active Pseudomonas Aeruginosa infection downregulates the SLC9A3 Cystic Fibrosis modifier gene.

## Introduction

RNA-Sequencing (RNA-Seq) has been widely used to measure the transcriptome for differential gene expression (DE) analysis, to identify genes that correlate with biological features of interest, such as a disease state or exposure. RNA-Seq produces data in the form of integer read counts to measure gene expression. Similar to many types of genomic data, due to the complexity of read alignment and quantification (SEQC/MAQC-III Consortium, 2014; Reuter et al., 2015), even after careful design, pre-processing, and quality control procedures, the RNA-Seq data can exhibit large variation and modeling challenges, with samples containing zero or low counts, outliers and variance heterogeneity. The noisy read counts create challenges for accurate detection of differentially expressed genes.

Numerous statistical methods have been developed for DE analysis. The most popular DE toolkits by citation are edgeR (Robinson et al., 2010), DESeq2 (Love et al., 2014) and limma (Ritchie et al., 2015). These toolkits comprise several pipelines by using a combination of different normalization, variance modelling and testing methods. A primary focus of these methods has been to model the mean-variance relationship of the count data at a given gene and provide accurate inference in small sample sizes. Most of these methods rely on two assumptions: first, a quadratic mean-variance structure is assumed to capture the overdispersion in the RNA-Seq read count. Second, different genes with similar count size should share similar variance, enabling the “borrowing of information” for variance modelling to achieve more reliable performance for studies with small samples. In particular, the primary configuration in edgeR and DESeq2 assumes a negative binomial (NB) distribution for the read count, with empirical Bayes shrinkage (Robinson and Smyth, 2007; Love et al., 2014) for dispersion estimation. Limma uses a normal transformation for the count data and applies voom variance modelling (Law et al., 2014). The voom method provides a non-parametric LOWESS estimate of the gene level mean-variance trend on the log-count per million. The individual level variance is estimated by interpolating the trend by the predicted count, then inputting the estimated variance as an inverse weight to the limmatrend pipeline. As a result, the linear model previously applied for the analysis of DE with microarray data can now also be used for DE analysis on the transformed RNA-Seq read counts.

The specific form of a quadratic mean-variance relationship may, however, be too restrictive to represent the actual dispersion pattern genome-wide in an RNA-Seq dataset. Although empirical Bayes provides some degree of robustness against model violation, the required specification of hyperparameter and tuning parameters can be difficult. There are other tools that relax the quadratic mean-variance assumption; for example, tweeDEseq (Esnaola et al., 2013) which is built on a more general Poisson-Tweedie family of distributions, and SAMSeq (Li and Tibshirani, 2013) which used a nonparametric Wilcoxon test. But covariate adjustment is challenging for these tools: Poisson-Tweedie regression is computationally expensive and may not converge (Bonat et al., 2018), while the permutation methods are not trivial for multiple explanatory variables.

Due to the complex nature of RNA-Seq data, several comparative studies have shown, even in well-powered studies, that different tools tend to detect different DE genes in the same dataset (Assefa et al., 2018). More-over, no single DE tool is uniformly superior to the other methods and the performance of all tools is strongly affected by sample size and biological variability (Conesa et al., 2016).

With advancements in sequencing technology and corresponding reductions in cost, RNA-Seq experiments with larger sample sizes are becoming more common-place (Wilks et al., 2021). Therefore, a paradigm shift for DE analysis that implements robust tools that rely on large sample statistical properties is now possible. Here we present Robust Profile likelihood-based differential Expression analysis (RoPE), a method for DE analysis that avoids the assumptions historically necessary to account for small sample sizes. Our method is inspired by the theory of robust profile likelihoods for mis-specified models (Royall and Tsou, 2003) and its application to generalized linear models (Blume et al., 2007). The important insight here is to treat dispersion as a model misspecification problem and make a robust adjustment to the likelihood function empirically from the data at a given gene. This is in contrast to methods such as DESeq2 and edgeR that specify a form of variance function genome-wide that may or may not hold at a given gene and plug in an estimate of the dispersion level based on that assumed relationship. We show that asymptotic tests based on RoPE provide reliable inference. We demonstrate the asymptotic properties of RoPE and its performance, compared with existing methods, through parametric simulation and non-parametric simulation studies. The diverse set of simulation studies show that RoPE is consistently a top-performer in large samples, in terms of sensitivity and error rate control.

We apply RoPE in a DE analysis using RNA-Seq of naive human nasal epithelia (HNE) from individuals with Cystic Fibrosis (CF). CF affects multiple organs with complex genetic epidemiology that includes contributions from genes beyond the causal CFTR variants. The majority of morbidity and mortality in CF is due to lung disease, the sequalae of cycles of infection with pathogens such as Pseudomonas Aeruginosa (PsA) and inflammation. The largest genome-wide association study (GWAS) of lung disease in CF from the International CF Gene Modifier Consortium (n=6365) identified a locus at Chr5p providing the smallest genome-wide significant p-values (Corvol et al., 2015). This locus contains a previously implicated CF modifier gene SLC9A3, reported to contribute to susceptibility to PsA infection (Dorfman et al., 2011), but the associated variants do not tag protein coding variation suggesting the causal variation impacts gene regulation. We hypothesize that variation in expression of SLC9A3 influences lung function in individuals with CF and that this relationship is mediated by bacterial infection with PsA. Hence, together with existing DE methods, we use RoPE to examine how the expression of SLC9A3 in human nasal epithelia, an airway model for CF lung disease (Knowles et al., 1981), is impacted in CF individuals with and without an active PsA infection.

## Methods

### Poisson log-linear model as a working model

We propose RoPE to detect gene expression differences using robust profile likelihood ratio test statistics. Similar to other widely implemented DE tools (McCarthy et al., 2012; Love et al., 2014), we apply a generalized linear model (GLM) for the RNA-Seq read counts. The GLM log-linear model is flexible to enable adjustment for technical variability such as sequencing depth, batch effects, sample quality, and it can also accommodate complex study designs. For a typical RNA-seq data matrix *Y* with *G* genes and *n* samples, the entries *y*_*gi*_ are the numbers of sequencing reads mapped to the *g*th gene in sample *i*. We choose the Poisson distribution as a working model, so *y*_*gi*_ ∼ Poisson 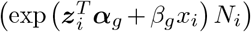 where the parameter of interest *β*_*g*_ is the fold change of gene *g* with respect to the experimental condition *x*_*i*_. Gene level intercepts and other sample-specific covariates are captured by 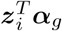. The trimmed mean of M-values (TMM) normalization method (Robinson and Oshlack, 2010) calculates scaling factors for each sample and multiplies by the total sample read count, *N*_*i*_, thus the relative abundance can be compared across samples regardless of the library size differences between the samples. Let 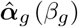 be the constrained parameter value maximized with respect to *β*_*g*_ and the scaled normalized library size 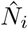 acts as an offset term in the GLM model. Then the corresponding gene level profile likelihood for the working model is

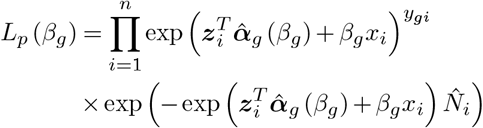

Implementation of profile likelihood function computation is motivated by the algorithm from the R package ProfileLikelihood (Choi, 2022). At a fixed *β*_*g*_ value, the profile likelihood function *L*_*p*_ (*β*_*g*_) is the likelihood function evaluated at the maximum likelihood estimator (MLE) for the Poisson GLM, providing *β*_*g*_*x*_*i*_ as the offset term. We apply the `mglmLevenberg` function from edgeR (McCarthy et al., 2012) with dispersion = 0 for estimation of the working model.

### Treating dispersion as model misspecification

RNA-seq read data is often over-dispersed, and when this over-dispersion arises the Poisson model fails to represent the extra variation. Instead of implementing a different parametric form for all genes analyzed in anticipation of the over-dispersion, such as negative binomial or a type of Tweedie distribution, we choose the Poisson regression model as a working model. We use a data-driven robust adjustment to the profile likelihood function, *L*_*p*_ (*β*_*g*_), to adjust for departures from the Poisson mean-variance assumption. The premise of this proposed approach is that even if the model is misspecified, the MLE, 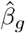,from the working model would still converge to the fold change estimate *β*_*g*_ from the unknown true model. This convergence result holds despite the standard error around the estimator being distorted, due to the violation of Bartlett’s second identity. This premise is satisfied since a canonical log link function was used in the GLM for the exponential family data and the interpretation of the regression parameter estimate remains unchanged by using the working model instead of the true model (Blume et al., 2007).

### Gene-level robust adjustment

We aim to correct the dispersion to the profile likelihood of gene expression change by following the procedure of parameter partitions of information matrices under the true model. The adjustment factors are defined by *A*(*α*_*g*_, *β*_*g*_), the inverse of observed information for the parameter of interest, and *B*(*α*_*g*_, *β*_*g*_), the inverse of the Godambe information (Godambe, 1960). For the purpose of restoring Bartlett’s second identity, the robust profile likelihood defined as 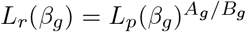 can achieve the correct information matrix under the unknown true model in large samples. In application, we implement the robust adjustment empirically. First, we define the predicted ex-pression count as 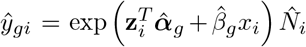, and residuals as 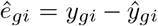. Then adjustment terms can be estimated by their asymptotically equivalent quantities as 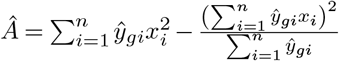 and 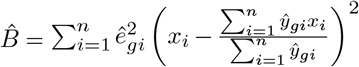. The resulting robust profile likelihood can be used to derive the standard large sample tests with the correct variance under the unknown true model, regardless of the exact meanvariance structure.

### Hypothesis testing for differential expression

For each gene, a working Poisson regression model is fitted: 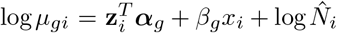. Given the dimension of the parameter of interest is 1, we have the asymptotic result of the (log-)robust profile likelihood ratio provides a test statistic (Stafford, 1996) 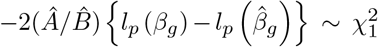. The question of differential expression is translated into assessing the null hypothesis of *β*_*g*_ = 0. Gene-wise *p*-values are adjusted by the Benjamini and Hochberg (1995) method (BH) to account for multiple hypothesis testing.

For a standard DE analysis, RoPE calculates the adjusted profile likelihood at MLE 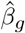 and at null 0. Additionally, RoPE can compute the profile likelihood *L*_*p*_(*β*_*g*_) and adjusted profile likelihood *L*_*r*_(*β*_*g*_) over values in the neighbourhood of 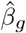 by an auto grid search algorithm as illustrated in figure 4. The standardized (relative to MLE) likelihoods over the grid can be utilized to find likelihood intervals that are analogous to confidence intervals and can serve as an alternative measure of inference (Pawitan, 2001; Strug, 2018).

## Simulation studies

We conducted extensive simulation studies to illustrate the performance of RoPE. RoPE was compared with the most frequently implemented DE tools: edgeR (Robinson et al., 2010) (default LRT option (McCarthy et al., 2012) and robust option (Zhou et al., 2014)), DESeq2 (Love et al., 2014) and limma-voom (Law et al., 2014). The selected DE tools, in contrast to other published tools (eg tweeDEseq, SAMSeq), run a fast integrated pipeline from normalization to testing, with underlying regression models accommodating covariates and complex designs. As all performance metrics depend on the statistical significance level, a reliable DE tool should demonstrate high sensitivity and statistical power at a controlled false discovery rate (Figures 1 and 3); and the actual false discovery rate should be similar to the nominal level (Figure 2). These main metrics are presented for the parametric and non-parametric simulation results.

**Figure 1.**
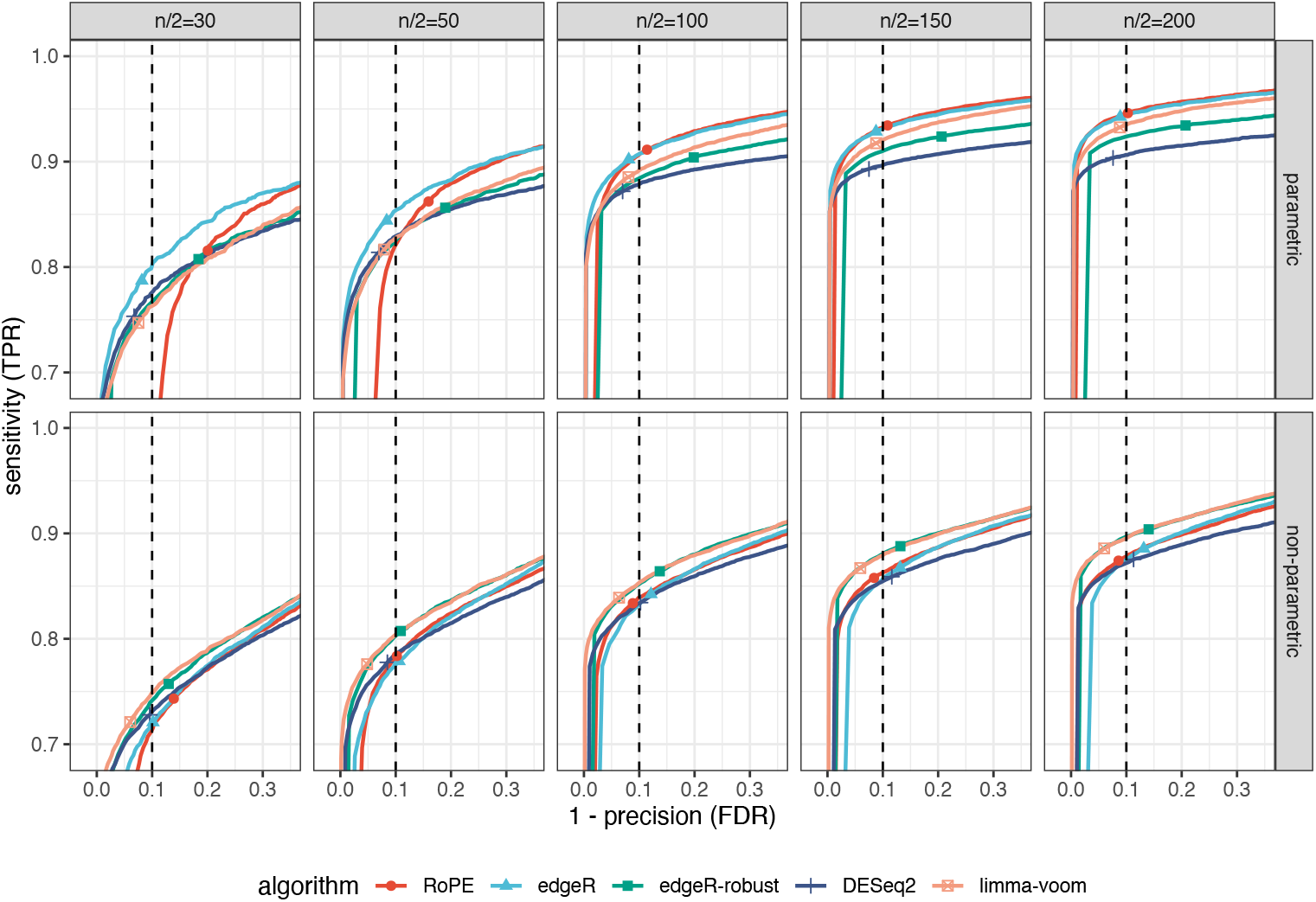
Simulation result: TPR vs FDR. Sensitivity (true positive rate) and 1 − precision (false discovery rate) of DE tools from the simulation study of two group comparisons at group sizes of 30, 50, 100, 150 and 200. RoPE exhibits substantial improvement of performance as sample sizes increases, and it shows high sensitivity over the range of false discovery rates for group sizes greater than 100 under the parametric simulation. Voom and edgeR-robust have high sensitivity over the range of false discovery rates under the non-parametric simulation. The points on the curve indicate the actual sensitivity and false discovery rate at the 0.1 adjusted *p*-value cutoff.

**Figure 2.**
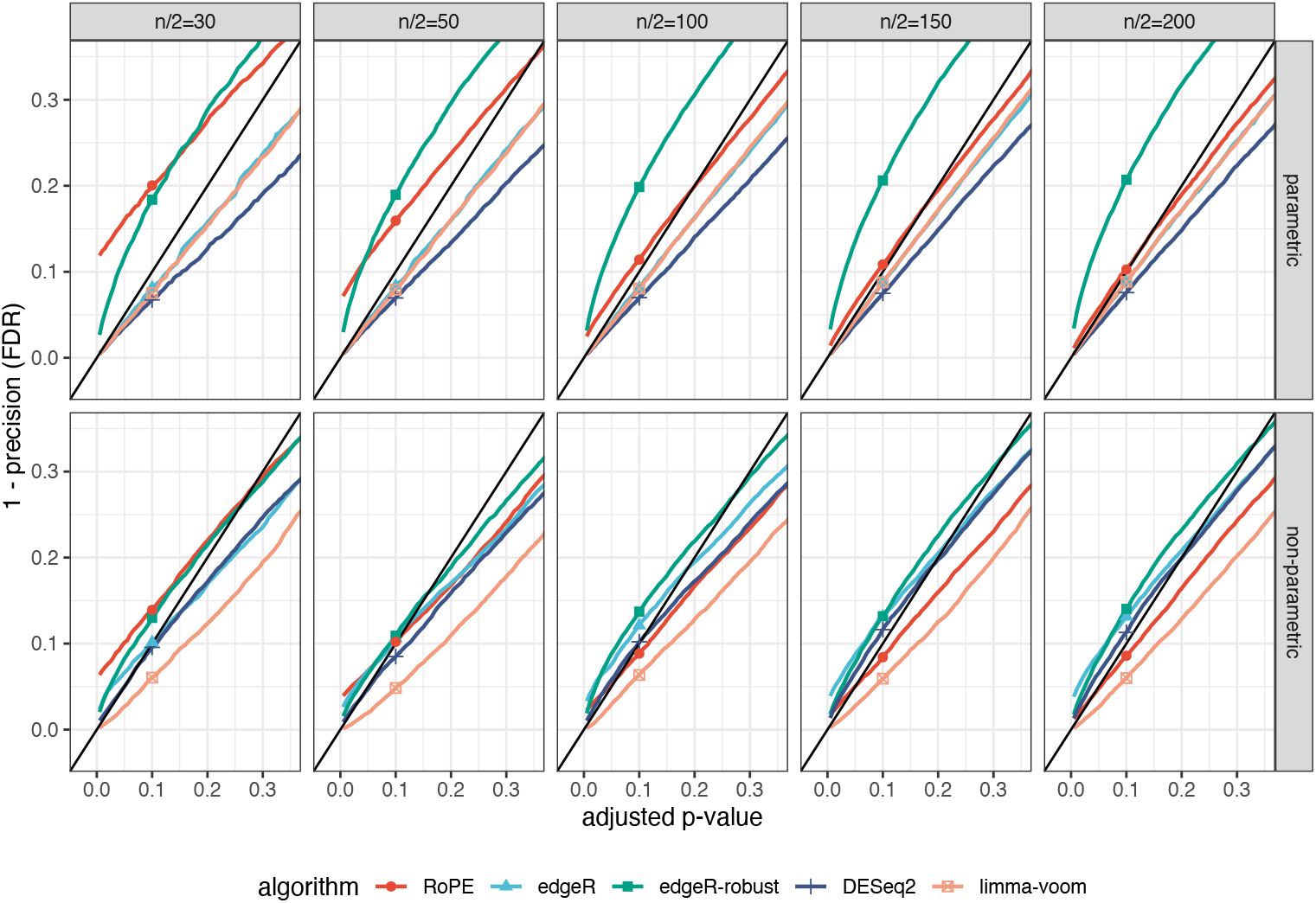
Simulation result: FDR vs adjusted-pvalue. 1 −precision (false discovery rate) and adjusted *p*-value threshold from the simulation study of two group comparisons at group sizes of 30, 50, 100, 150 and 200. When the group sizes reach 100, the alignment of RoPE with the 45-degree line reflects proper control of the false discovery rate under both parametric and non-parametric simulation settings.

**Figure 3.**
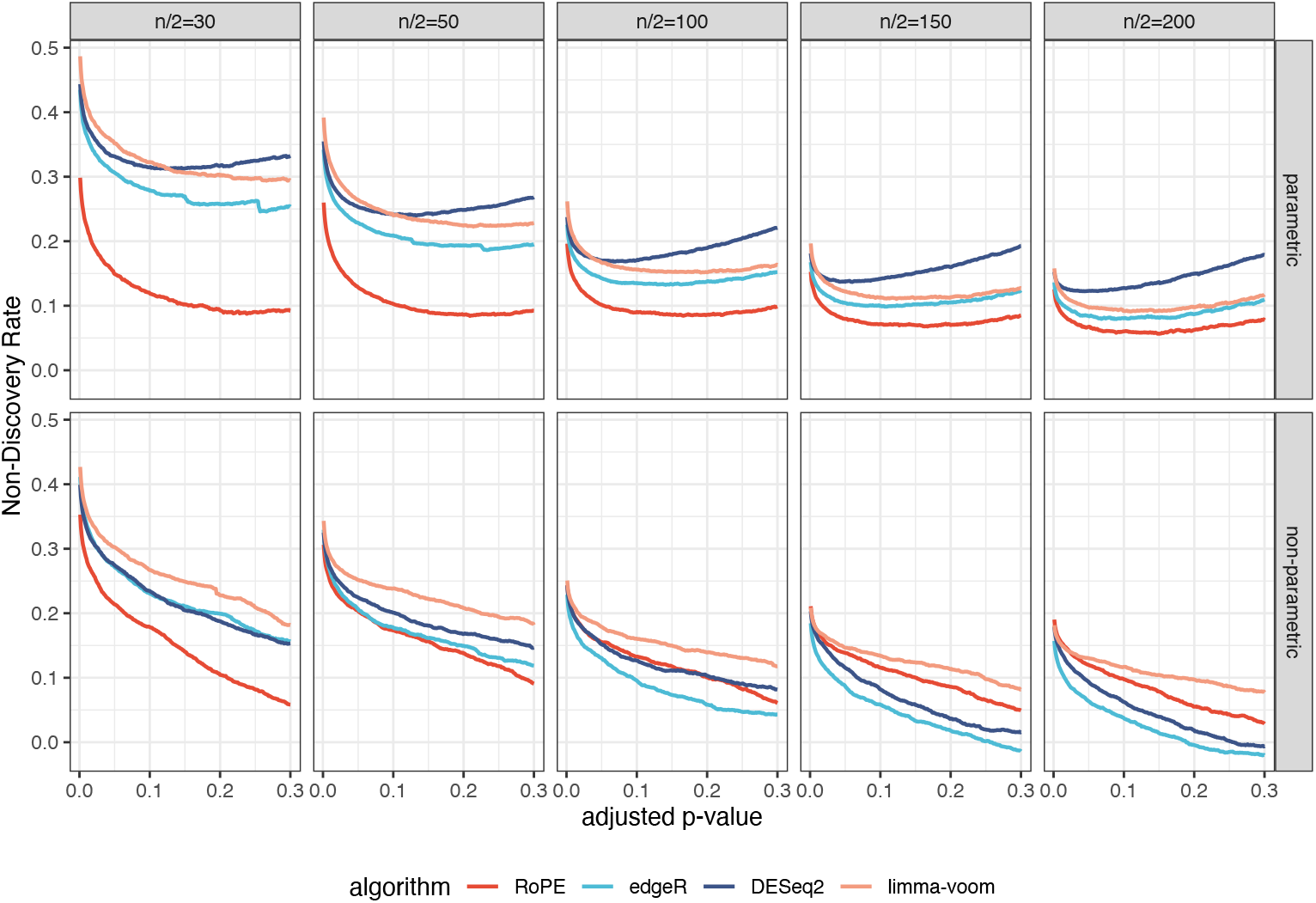
Simulation result: NDR vs adjusted-pvalue. 1 −power (non-discovery rate) and adjusted *p*-value threshold from the simulation study of two group comparisons at group sizes of 30, 50, 100, 150 and 200. RoPE displays low NDR, and other methods show decreasing NDR over increasing sample sizes. edgeR-robust was removed from this comparison due to the biased NDR values resulting from the severely inflated error rate indicated by Figure 2. When the true power approaches 1, the large variation from simulated non-parametric replicates pushes the median NDR measure slightly below 0.

### Simulation design

For a comprehensive unbiased assessment, we model the simulation studies after Love et al. (2014) for the parametric negative binomial simulation, and from Assefa et al. (2018) for the non-parametric sub-sampling simulation. The RNA-Seq datasets were simulated for a two-group comparison under varying equal group sizes (*n/*2 = 30, 50, 100, 150, 200). At each sample size level, 50 datasets of 15000 genes (*G*) were simulated, with 20% of the genes being truly differentially expressed. Genes with zero counts for all samples were removed, then the detection of DE genes is determined by the BH adjusted *p*-values from each method on the non-zero samples. Since in multiple testing, the false discovery rate is a widely used alternative measure of the type I error. To measure the type II error rate for multiple comparisons, we hereby apply the non-discovery rate (Craiu and Sun, 2008), defined by the expected proportion of false negatives among the true alternative hypotheses. For each dataset, sensitivity is calculated as the proportion of significant DE genes among the true DE genes, false positive rate (1− specificity) is the proportion of significant DE genes among the true non-DE genes, non-discovery rate (1− power) is the proportion of estimated non-significant genes among the true DE genes, and false discovery rate (1− precision) is the proportion of true non-DE genes among the significant genes. The benchmark metrics were computed as the median of corresponding actual metric measures from 50 replicates at different adjusted *p*-value thresholds.

#### Parametric simulations

We generate RNA-Seq counts from the negative binomial distribution by sampling 4000 mean and dispersion level pairs from an RNASeq dataset (Pickrell et al., 2010) (Gene Expression Omnibus: GSE19480) of 69 lymphoblastoid cell lines from unrelated Nigerian individuals. At each sample size, 80% of the non-DE genes are set to have the same mean parameter for both groups, and the 20% DE genes have mean differences between groups by a fold change of 2 or 0.5 with equal chance.

#### Non-parametric simulations

We use SimSeq (Benidt and Nettleton, 2015) to simulate RNA-Seq counts non-parametrically by sub-sampling the Esophagus Mucosa (*n*_1_ = 407) and the Lung (*n*_2_ = 427) tissues of protein coding genes from the Genotype-Tissue Expression project (GTEX) (Lonsdale et al., 2013) v7. The 20% DE genes are randomly selected by the probability sampling weights, where the weights are determined by the local fdr from a Wilcoxon DE test, and 80% non-DE genes are randomly selected by equal weights from the rest of the genes. Then, simulated datasets are constructed by sampling without replacement within each tissue group for the selected DE and non-DE genes, until the pre-specified sample sizes are achieved.

### Simulation results

Under the parametric simulation, RoPE shows an inflated false discovery rate at group sizes of 30 and 50 compared to the other tools. As group sizes increase to 100, RoPE demonstrates high sensitivity while balancing precision at a controlled false discovery rate. Whereas edgeR shows similar performance to RoPE, edgeR-robust has inflated false discovery rate even with large sample sizes. DESeq2 and voom appear to be conservative in the DE significance conclusions, resulting in the actual false discovery rate lower than the nominal adjusted *p*-values, and a lower sensitivity but slightly higher specificity and precision.

As the GTEX samples contain larger biological variability, and non-parametric simulated datasets are purported to reproduce real-world mean-variance complexity (Benidt and Nettleton, 2015), all methods have decreased sensitivity compared to the negative binomial simulation. Under the non-parametric simulation for group sizes greater than 100, RoPE demonstrates satisfactory sensitivity and specificity, while keeping the best control of false discovery rate as in the NB simulation. In contrast, edgeR and DESeq2 fail to control the false discovery rate at the same level as in the NB simulation. The voom method has the highest sensitivity and the lowest false discovery rate, but voom also results in the lowest number of significant genes. Overall, RoPE exhibits competitive performance with a controlled false discovery rate in all large sample settings and for the different simulation models implemented in the literature. It is also of note that RoPE has the highest proportion of rejections (Supplementary file: Figure S1) in small samples, resulting in the highest statistical power (equivalently the lowest non-discovery rate).

We also investigate the trade-off between sensitivity and false positive rates, as well as the impact of varying the percentage of true DE gene from 5% to 30% (Supplementary file: Figures S2-S7). The simulated samples with a lower DE percentage seem to result in an inflated false discovery rate for RoPE in smaller samples, but RoPE displays higher sensitivity and lower false positive rates in samples with a high DE percentage. We also verify the robust LR test statistics’ distributional properties by showing the convergence to the theoretical quantile under the null hypothesis as the sample size increases through the chi-square quantile-quantile plot (Supplementary file: Figures S7). With group sizes greater than 150, the Q-Q plots are below the 45 degrees line, implying better false-positive control under the null hypothesis.

## Differential Gene Expression Analysis of PsA Infection in CF human nasal epithelia

We conducted differential gene expression analysis with RNA-Seq of RNA from human nasal epithelial (HNE) cells obtained from 65 CF donors enrolled in the CF Canada SickKids program in individualized CF therapy (Polineni et al., 2018; Eckford et al., 2019). The nasal epithelia is a surrogate airway model in CF (Keegan and Brewington, 2021), where the PsA infection status was identified from upper or lower respiratory tract cultures. The sample quality control implemented ensures the individuals are unrelated, with less than 2 weeks between PsA measurement and nasal sample and sample quality measured by the RNA integrity number greater than 7. Low count genes that have more than 30% of the samples per group with zero counts were removed, leaving 29330 genes for DE analysis after filtering.

We apply RoPE for DE analysis to examine whether there is differential expression of SLC9A3 between *n* = 27 samples who were infected with the common CF pathogen Pseudomonas Aeruginosa at the time of nasal scrape and n=38 individuals who were without an active PsA infection. The sample characteristics are described in Table 1. For exploratory analysis and method evaluation, we assess DE genome-wide. We fitted RoPE for each gene and included the genotype PCs, study site, RNA integrity number, sex and CD45 expression as a marker of immune cell proportion, as covariates in the model. The same pipeline was implemented for edgeR, DESeq2 and limma-voom for comparison.

**Table 1.** Characteristics of the human nasal epithelia donors and RNA undergoing RNA-Seq.

|  | Level | Overall |
| --- | --- | --- |
| n |  | 65 |
| Sex (%) | Male | 35 (53.8) |
|  | Female | 30 (46.2) |
| RNA integrity number (mean (SD)) |  | 8.92 (0.81) |
| immune cell proportion (mean (SD)) |  | 0.21 (0.06) |
| PsA Status at brush (%) | Grown in sputum | 27 (41.5) |
|  | No Growth | 38 (58.5) |

SLC9A3 showed strong evidence of DE using RoPE (*p*-value = 2.45 × 10^−6^; Table 2, Figure 4). All other methods support the association (edgeR: *p*-value= 3.75 ×10^−5^, DESeq2: *p*-value= 2.98 ×10^−5^ voom: *p*-value= 5.12 ×10^−3^), which suggests active PsA infection appears to downregulate the expression of SLC9A3 in individuals with CF, although only RoPE provides genome-wide significant evidence.

**Table 2.** Summary of differential expression analysis of SLC9A3. For each method, this summary table contains the estimates of natural logarithm of differential expression fold changes (logFC), the nominal *p*-values, the BH adjusted *p*-values (Adj *p*-value) for DE tests genome-wide, and the rank of DE significance by adjusted *p*-values for SLC9A3.

| Method | logFC | *p*-value | BH Adj *p*-value | Rank |
| --- | --- | --- | --- | --- |
| edgeR | -1.68 | $3.75 \times 10^{-5}$ | 0.1226 | 8 |
| DESeq2 | -1.16 | $2.98 \times 10^{-5}$ | 0.0972 | 9 |
| voom | -1.39 | $5.12 \times 10^{-3}$ | 0.9998 | 92 |
| RoPE | -1.27 | $2.45 \times 10^{-6}$ | 0.0041 | 16 |

**Figure 4.**
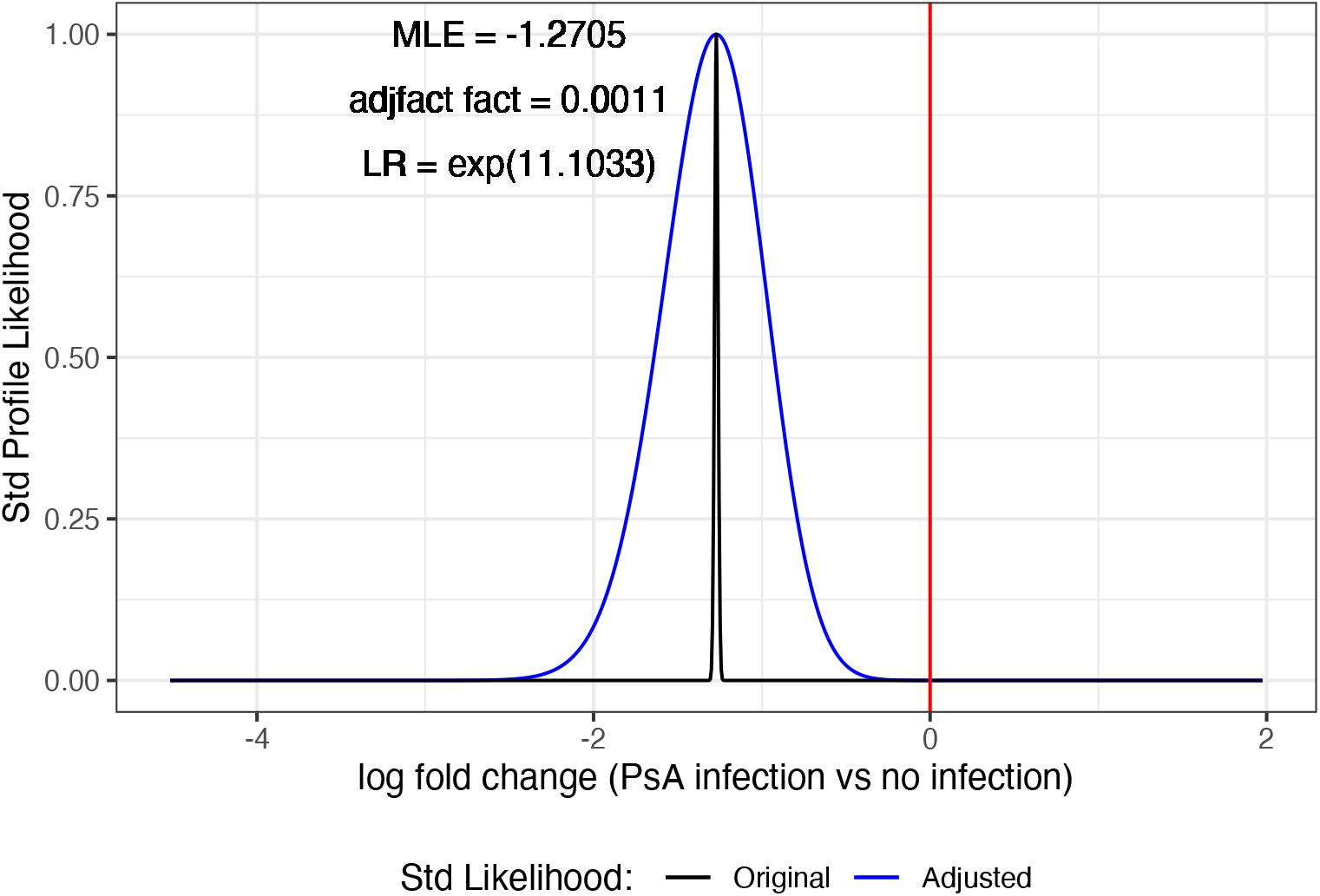
Standardized Profile Likelihood for SLC9A3. Standardized profile likelihood of log fold change before and after robust adjustment for SLC9A3 from DE analysis of PsA infection. The profile MLE is the log fold change of −1.27 (−1.83 in log 2*FC*). The robust profile likelihood ratio (LR) is *e*^11.1033^. This gene is significantly DE, even after using the robust adjustment to account for the overdispersion in the data.

In genome-wide analysis, RoPE detected 20 genes up-regulated in those with active PsA infection and 39 genes downregulated at the 0.05 FDR level (Figure 5, Supplementary file: Table S2, Figure S9). For comparison, edgeR, DESeq2 and voom detected 4, 4, and 0 DE genes, respectively at the 0.05 FDR level). Examining the similarity in results between DE methods, the top 20 DE genes from RoPE, edgeR and DESeq2 have 6 genes in common (Figure 6), including SLC9A3 (Table 2). Notably, however, only RoPE generated a genome-wide adjusted *p*-value of less than 0.05 for SLC9A3. Among the genes showing DE at the adjusted *p*-value level of *<* 0.05 with RoPE (Supplementary file: Table S2), SLC9A3 (Dorfman et al., 2011) and SLC26A4 (Kim et al., 2019; Rehman et al., 2020) (Supplementary file: Table S1) have been previously implicated in CF infection phenotypes.

**Figure 5.**
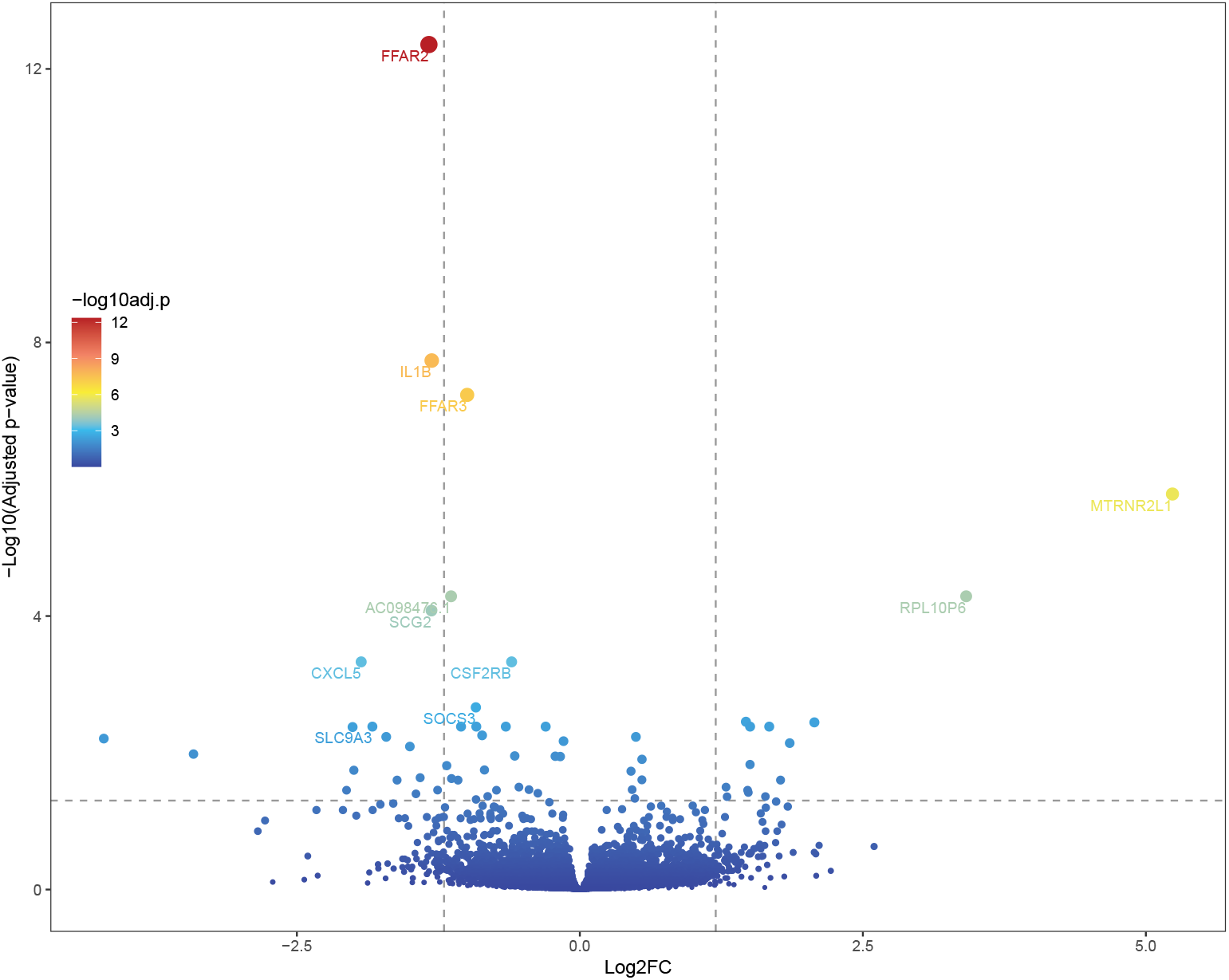
Volcano plot contrasting log-fold change with adjusted p-values from the analysis of HNE samples with RoPE method. RoPE detected 59 genes at the 0.05 FDR level (above horizontal line) with top 10 signals and SLC9A3 are labelled. The vertical lines are effect size threshold of log 2*FC* = ±1.2.

**Figure 6.**
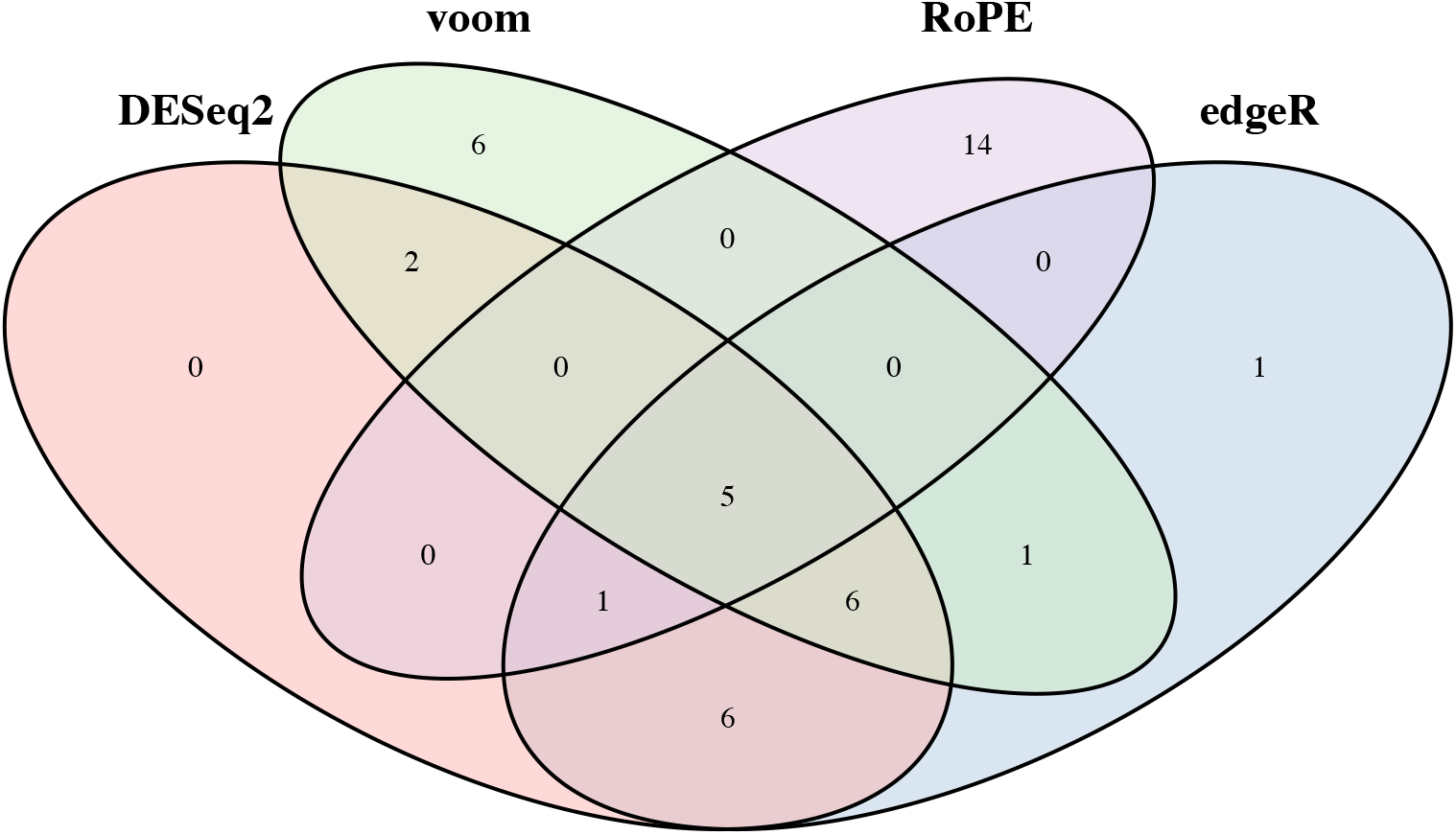
Venn diagram of the top 20 ranked DE genes. Resulting from application of edgeR, DESeq2, voom and RoPE on the HNE samples. The five common DE genes are: SCG2, AC098476.1, MRAP2, AC022826.1 and ZNF385D. Voom is the only method that did not rank SLC9A3 in its top 20.

Given the major difference in the number of genes returned as DE between RoPE and the other methods, and that the simulation studies indicated one should expect some false discovery rate inflation with RoPE at this sample size but greatest power, we conducted a pathway analysis on the GO Annotation (Ashburner et al., 2000; The Gene Ontology Consortium, 2020) to determine whether there was signal in the results. We inputted the 59 DE genes identified by RoPE to the gene ontology analysis to check relevant gene set clusters with the three GO databases: Biology process, cellular component and molecular function (Table 3). Supporting the hypothesis that RoPE is detecting truly differentially expressed genes, we identified pathways that would be expected given the infection phenotype: the inflammatory response (GO:0006954) and cytokine activity pathways (GO:0005125) were shown to be significant (*FDR <* 0.05). Two of the most significant genes genome-wide were the Free Fatty Acid Receptor 2 and 3 (FFAR2 and FFAR3) which are reported to be involved in the inflammatory response (Alvarez-Curto and Milligan, 2016), and these also contributed to several significant gene sets. Interestingly, FFAR2 and FFAR3 were ranked first and third by RoPE, but their rank using the other tools (edgeR: 710, 30; DESeq2: 775, 210; voom: 80, 5349) were low. SLC9A3 and SLC26A4 appear together on pathways (*FDR <* 0.10) of brush border membrane (GO:0031526) and ion transport (GO:0006811), suggesting further evidence of relevance for the DE genes and CF.

**Table 3.** Pathway enriched by PsA infection in CF nasal epithelium. The 59 DE genes identified by RoPE were imputed to the gene ontology analysis. Eight pathways are shown to be significant with *FDR <* 0.05.

| Genes | Process name | *p*-value | BH Adj *p*-value |
| --- | --- | --- | --- |
| ACOD1; FFAR3; MEFV;<br>IL1B; SCG2; TACR1;<br>IL17A; PROK2; TNFRSF4;<br>PTGER4; CXCL5 | inflammatory response | $1.80 \times 10^{-8}$ | $9.97 \times 10^{-6}$ |
| ACOD1; SOCS3; MEFV;<br>SAA1; PTGER4 | negative regulation of inflammatory response | $9.70 \times 10^{-6}$ | 0.0027 |
| FFAR3; FFAR2; | positive regulation of acute inflammatory response to non-antigenic stimulus | $5.08 \times 10^{-5}$ | 0.0070 |
| FFAR3; FFAR2; | regulation of peptide hormone secretion | $5.08 \times 10^{-5}$ | 0.0070 |
| SAA1; TACR1; PROK2;<br>PTGER4; CXCR1 | positive regulation of cytosolic calcium ion concentration | $8.69 \times 10^{-5}$ | 0.0096 |
| FFAR3; FFAR2; | mucosal immune response | $2.35 \times 10^{-4}$ | 0.0198 |
| IL1B; SCG2; LTB;<br>IL17A; SCGB3A1 | cytokine activity | $2.51 \times 10^{-4}$ | 0.0198 |
| SOCS3; IL1B; SAA1;<br>CSF2RB; IL17A; IL17REL | cytokine-mediated signaling pathway | $2.89 \times 10^{-4}$ | 0.0200 |

## Discussion and conclusions

With increasing sample sizes for differential gene expression studies, RoPE provides a simple, robust solution to detect DE in noisy count data. RoPE achieves theoretical error rate control, relaxes the negative binomial model assumption and bypasses the time-consuming procedure of dispersion estimation. The GLM regression provides a flexible framework to incorporate complex designs, while the robust adjustment to its profile likelihood handles all functional forms of dispersion patterns that may arise. The computational time of RoPE is very efficient and can run on a typical laptop, thanks to the simple analytic form of robust profile likelihood and the numerically stable and fast mglmLevenberg function from edgeR (McCarthy et al., 2012).

The main limitation of RoPE stems from the sample size requirement necessary to achieve robust inference and maintained error rate control. According to our simulation, when the RNA-Seq sample size is less than 30 per group, RoPE is likely to be liberal in detecting DE genes so that it will retain high sensitivity but the false discovery rate will be inflated. This was consistent with what was observed in the analysis of DE in the CF HNE application, where RoPE concluded many DE genes, but these results appeared to contain true signal. In the presence of outliers, RoPE estimates can be biased at small sample size. Previous work (Assefa et al., 2018) has shown that if an RNA-Seq dataset contains a high degree of variability and small effect sizes, all existing methods will exhibit substandard performance. Nevertheless, in our simulations under these challenging scenarios, RoPE has the best improvement in performance with increasing sample size and would outperform existing methods in these large-sample scenarios.

Cycles of infection and inflammation in CF lead to chronic, progressive lung disease. The CF gene modifier study identified the SLC9A3 locus as a contributor to lung disease and the association colocalized with gene expression of SLC9A3 in the lungs using the GTEx resource (Soave et al., 2015; Corvol et al., 2015). eQTL analysis using the CF HNEs was inconclusive, however the results here suggest that this may have been complicated by the subset of participants who were infected with PsA at the time of nasal brush. The GTEx eQTL analysis indicates that a higher genetically determined SLC9A3 expression is associated with worse lung function.

This study of gene expression differences upon infection with PsA identified SLC9A3 as a differentially expressed gene, down-regulated upon lung bacterial infection in CF individuals. This was supported by RoPE and the other contemporary DE tools, most notable DESeq2 and edgeR. After accounting for the covariates, RoPE estimated that SLC9A3 gene expression was decreased 72% on average for PsA infected individuals versus those without PsA infection. Together these results, as well as a previous candidate gene study that demonstrated SLC9A3 genotype to be associated with an earlier acquisition of PsA, suggests that a high SLC9A3 expressing environment may create a preferred milieu for pseudomonas aeruginosa. This is supported by previous work demonstrating that the age of chronic PsA is influenced by genetic modifiers and has high heritability (Green et al., 2012). This evidence could point to therapeutic targets that could ameliorate the damaging effects of PsA infection on CF lung function (Strug et al., 2018).

EdgeR, DESeq2 and limma voom identified very few DE genes at the 5% FDR level, while RoPE identified fifty-nine. From the simulation result (Figures 1,2,3) for comparable sample sizes, we would expect RoPE to have high sensitivity and statistical power at a cost of slightly inflated FDR. Concern that RoPE identified all false discoveries was ameliorated by the pathway analysis that demonstrated the inflammatory response as the most enriched pathway, since the DE genes were identified for the phenotype of active versus no PsA infection. It is worth mentioning that we did adjust for immune cell contamination by including CD45 expression in the regression model. Whether this adjustment is effective or not, RoPE identified DE genes that belong to pathways relevant to the infection phenotype or relevant to an incomplete immune cell contamination adjustment. This result is consistent with a previous study (Ogilvie et al., 2011) demonstrating that the inflammatory response is the most significant difference in CF versus non-CF bronchial epithelium.

The computationally efficient RoPE method is especially suited to differential gene expression analysis in large samples, which complements currently available DE analysis toolkits that cater to small sample size study designs. We recommend RoPE for RNA-Seq datasets with more than 50 replicates per group. The robust adjustment is intuitive and powerful and provides a data-driven modification on the parameter of interest. As a future extension to RoPE, we plan to apply bartlett corrections (DiCiccio and Stern, 1994) to other modified profile likelihoods that have better approximations for smaller dependence samples (Zucker et al., 2000). This underlying robust regression framework is not limited to differential expression analysis, but may be useful in other large-scale omics studies, and in particular studies of DNA methylation where dispersion estimation is fundamental to adequate inference.

## Supplementary Information

Supplementary material has been submitted as a separate file.

### Software

The main results presented in this article are generated by R version 4.1.2 and Bioconductor (Gentleman et al., 2004) packages edgeR 3.36.0, DESeq2 1.34.0 and limma 3.50.0. Pathway analysis was performed by the command line tool GeneSCF (Subhash and Kanduri, 2016). The RoPE method is implemented in R package roper and is available at https://github.com/strug-hub/roper.

### Funding

Funding was provided by Cystic Fibrosis Foundation STRUG17PO; Canadian Institutes of Health Research (FRN-167282), Cystic Fibrosis Canada (2626) and the CFIT Program funded by the SickKids Foundation and CF Canada; Natural Sciences and Engineering Research Council of Canada (RGPIN-2015-03742). This work was also funded by the Government of Canada through Genome Canada (OGI-148) and supported by a grant from the Government of Ontario.The funders of the study play no role in study design, data collection and analysis, decision to publish or preparation of the manuscript. L.Z. is a trainee of the CANSSI Ontario STAGE (Strategic Training for Advanced Genetic Epidemiology) program at the University of Toronto.

## Acknowledgments

We thank the Cystic Fibrosis Canada-SickKids Program for Individualized Therapy for access to the RNA-sequencing of the nasal epithelia.

## Ethics approval and consent to participate

The Canadian Gene Modifier Study (CGMS) was approved by the Research Ethics Board of the Hospital for Sick Children (# 0020020214 from 2012-2019 and # 1000065760 from 2019-present) and all participating sub-sites. Written informed consent was obtained from all participants or parents/guardians/substitute decision makers prior to inclusion in the study. The CGMS is approved by the Research Ethics Board of the Hospital for Sick Children for the usage of public and external data.

## Availability of data and materials

The datasets used and/or analysed during the current study are available from the CF Canada-SickKids Program for Individualized Therapy Biobank at https://lab.research.sickkids.ca/cfit/cystic-fibrosis-patients-families-researchers. The roper R package and code used to analyze the data is available under a GPLv3 license on the GitHub repository for this paper https://github.com/strug-hub/roper.

