## Supplementary file for "RoPE: a robust profile likelihood method for differential gene expression analysis"

### Supplementary Figures and Tables

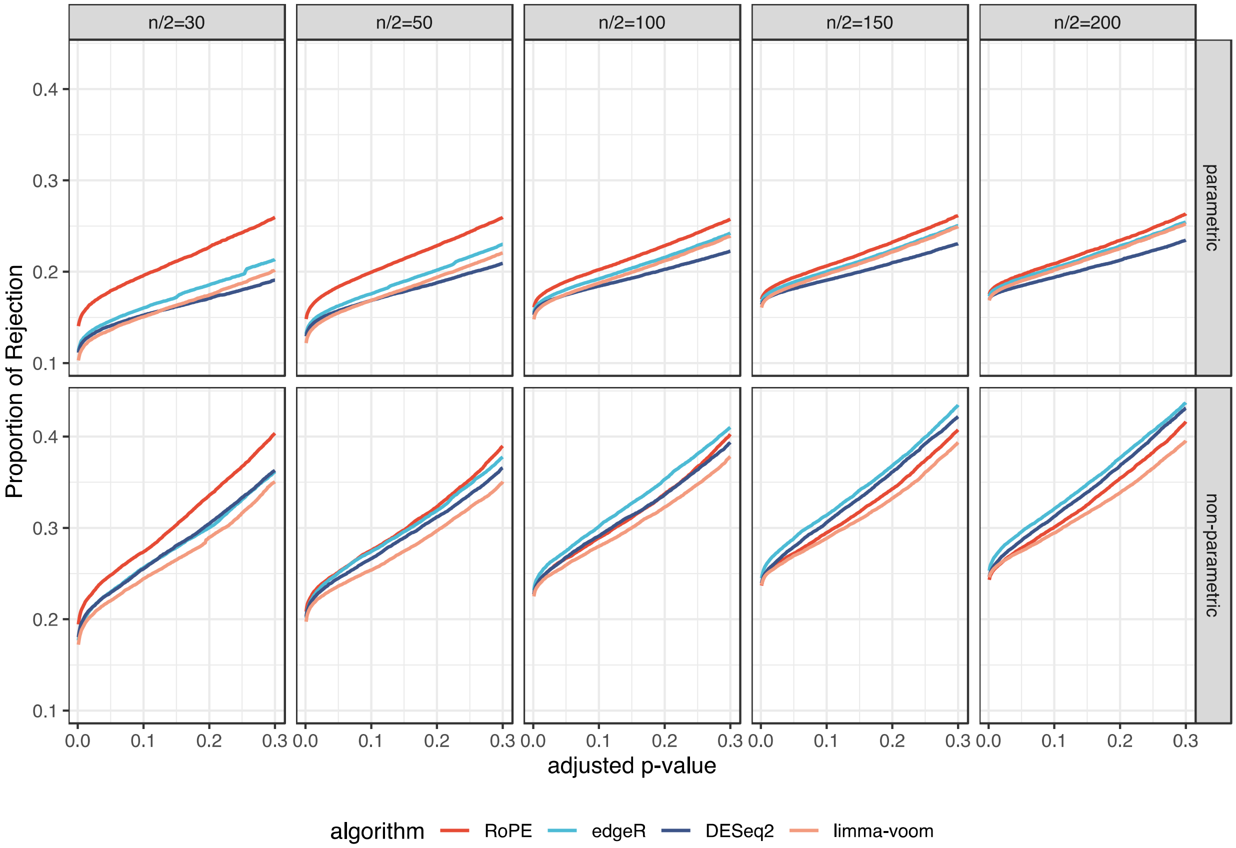


**Figure S1 True DE genes 20% simulation result: POR vs adjusted-pvalue.** Proportion of Rejection and adjusted p-value threshold from the simulation study of two group comparisons at group sizes of 30, 50, 100, 150 and 200.

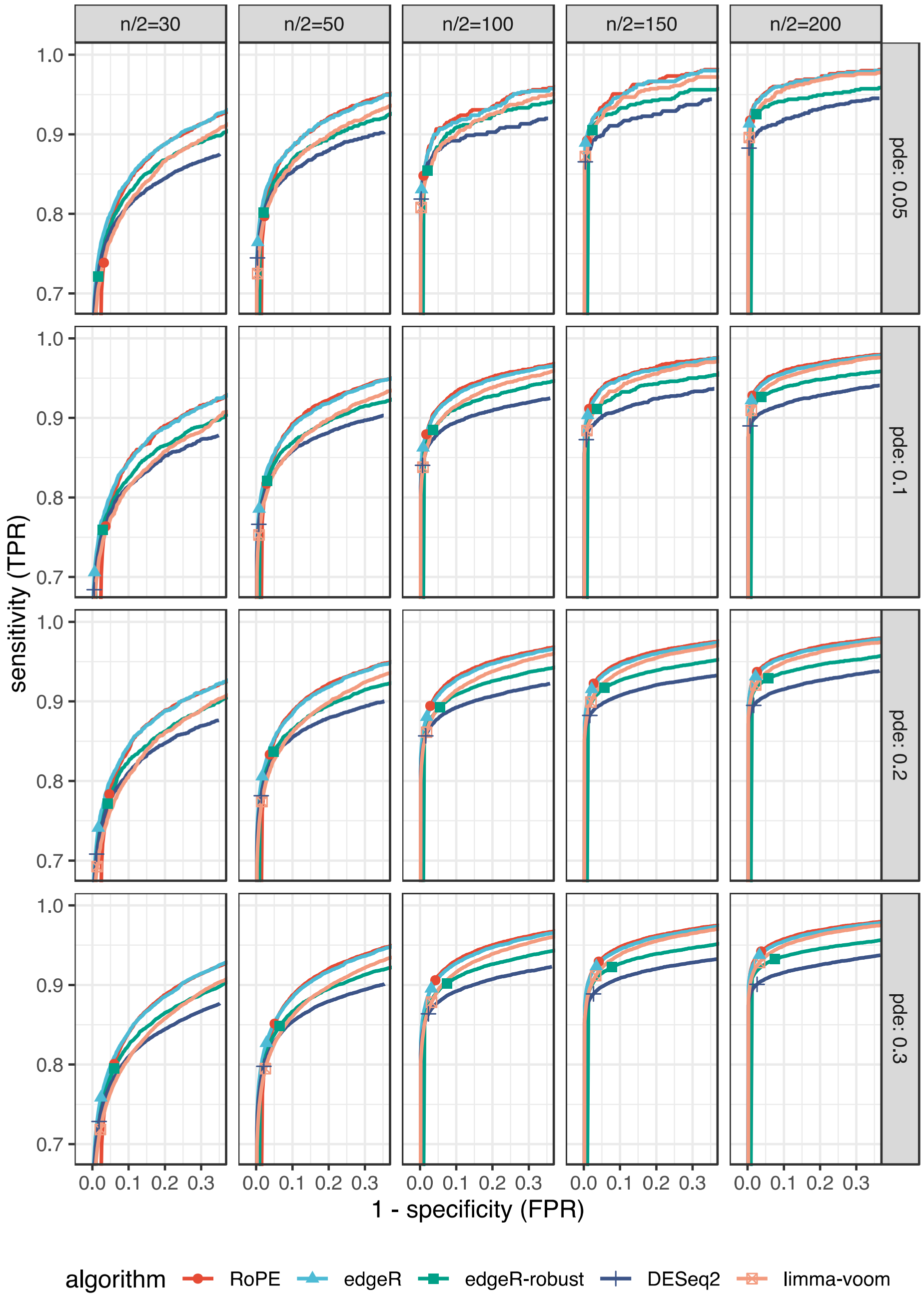


**Figure S2: Parametric simulation result varying the percentage of true DE gene from 5% to 30%: TPR vs FPR**

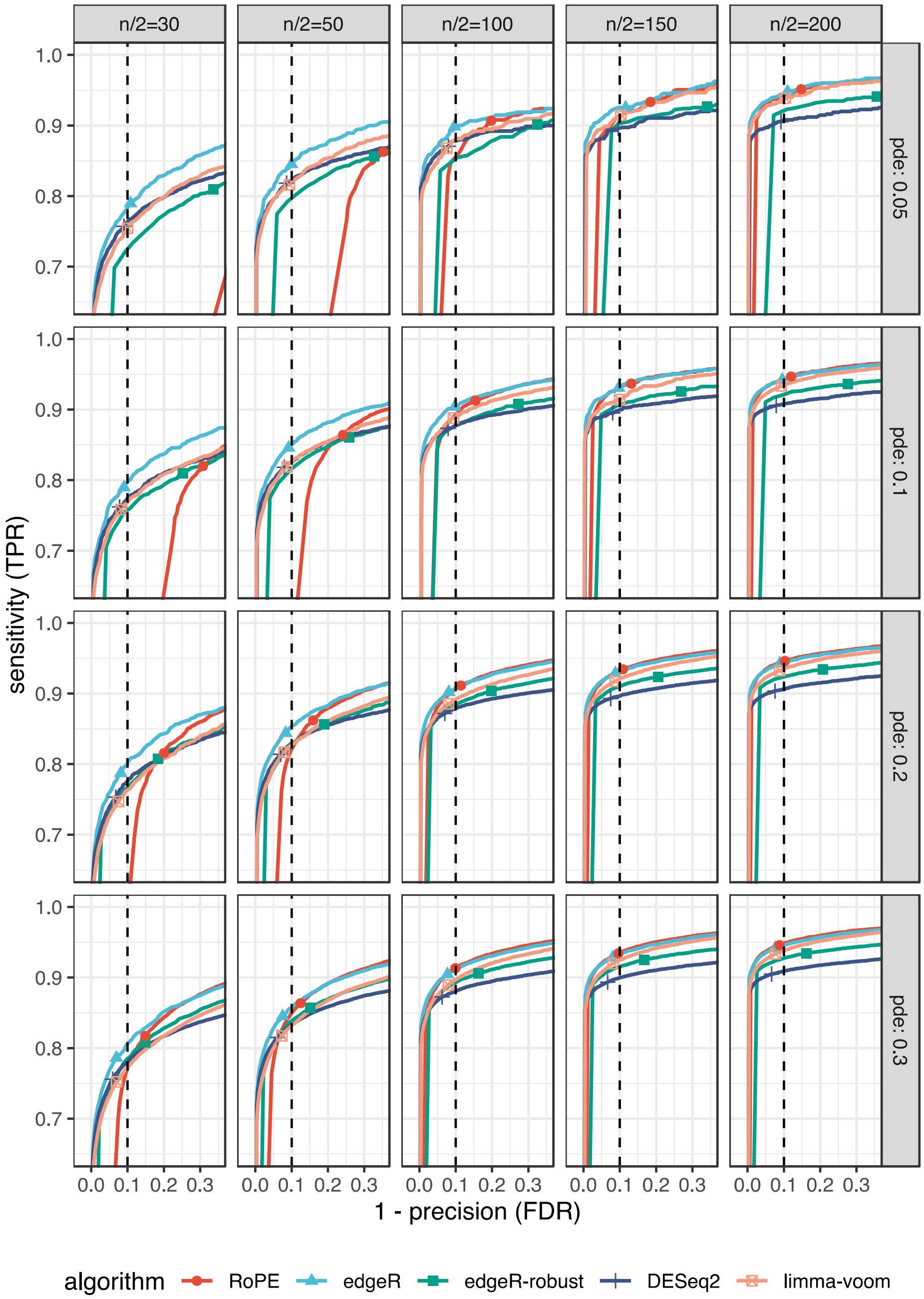


**Figure S3: Parametric simulation result varying the percentage of true DE gene from 5% to 30%: TPR vs FDR**

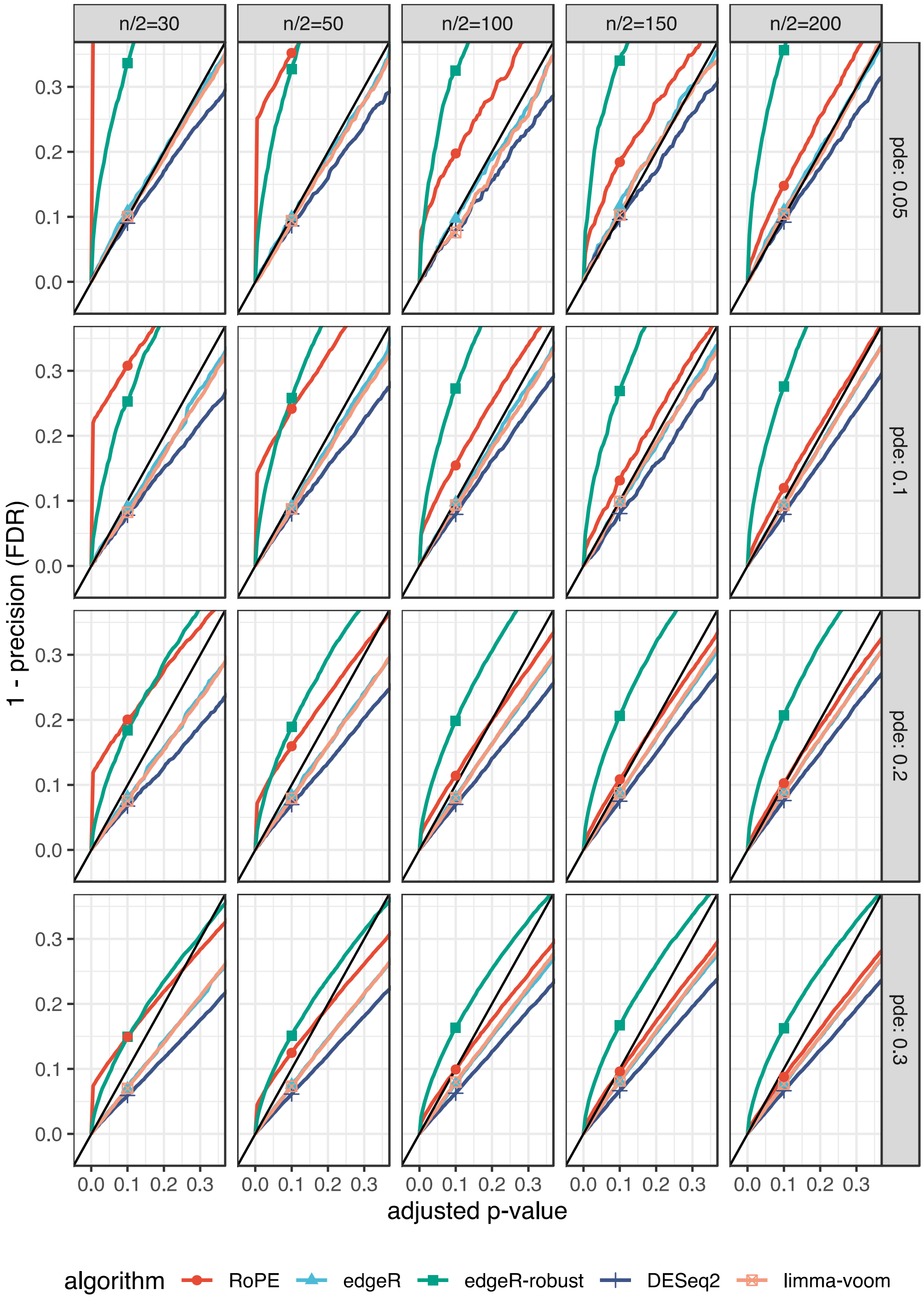


**Figure S4: Parametric simulation result varying the percentage of true DE gene from 5% to 30%: FDR vs Adj-Pvalue**

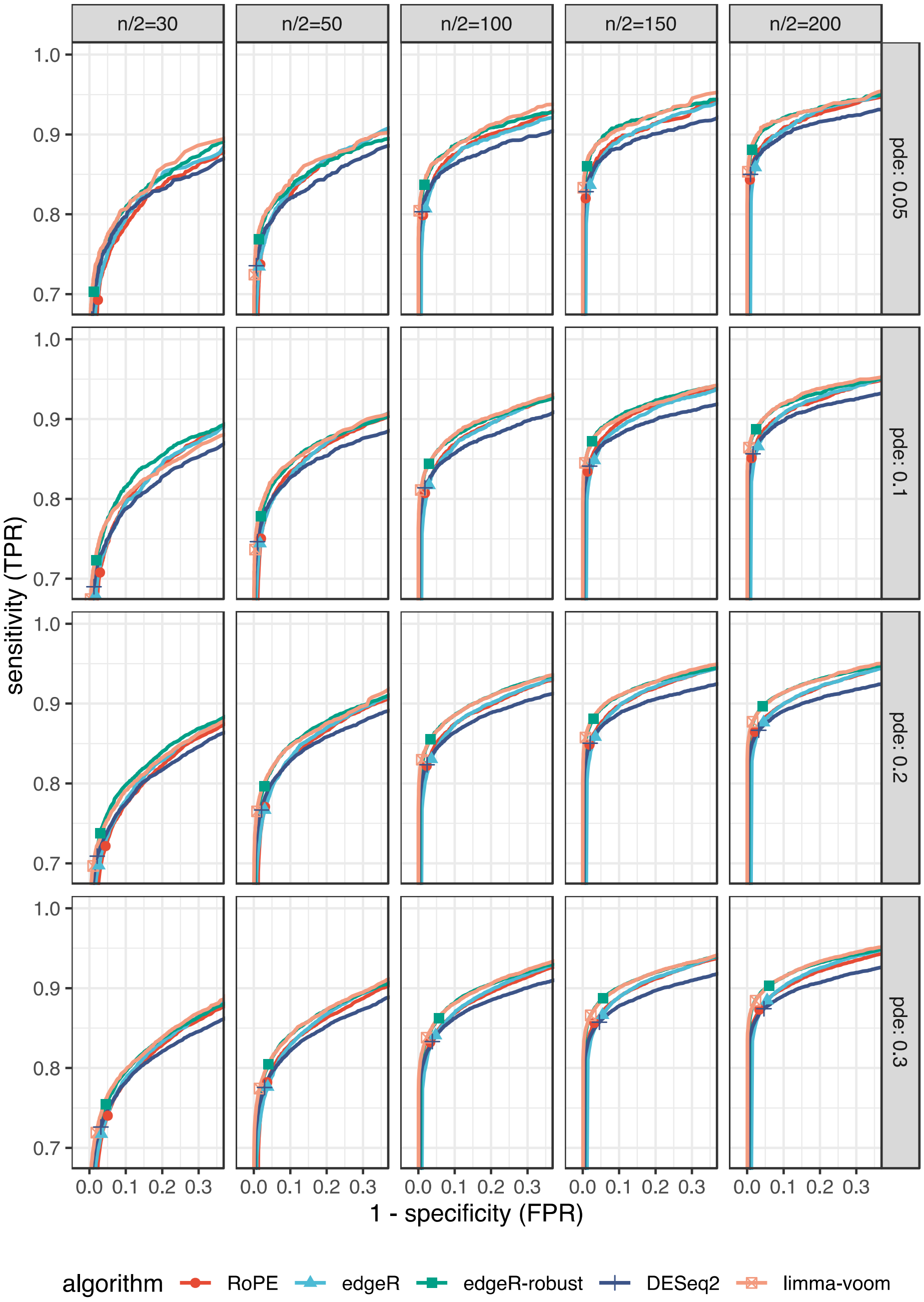


**Figure S5: Non-parametric simulation result varying the percentage of true DE gene from 5% to 30%: TPR vs FPR**

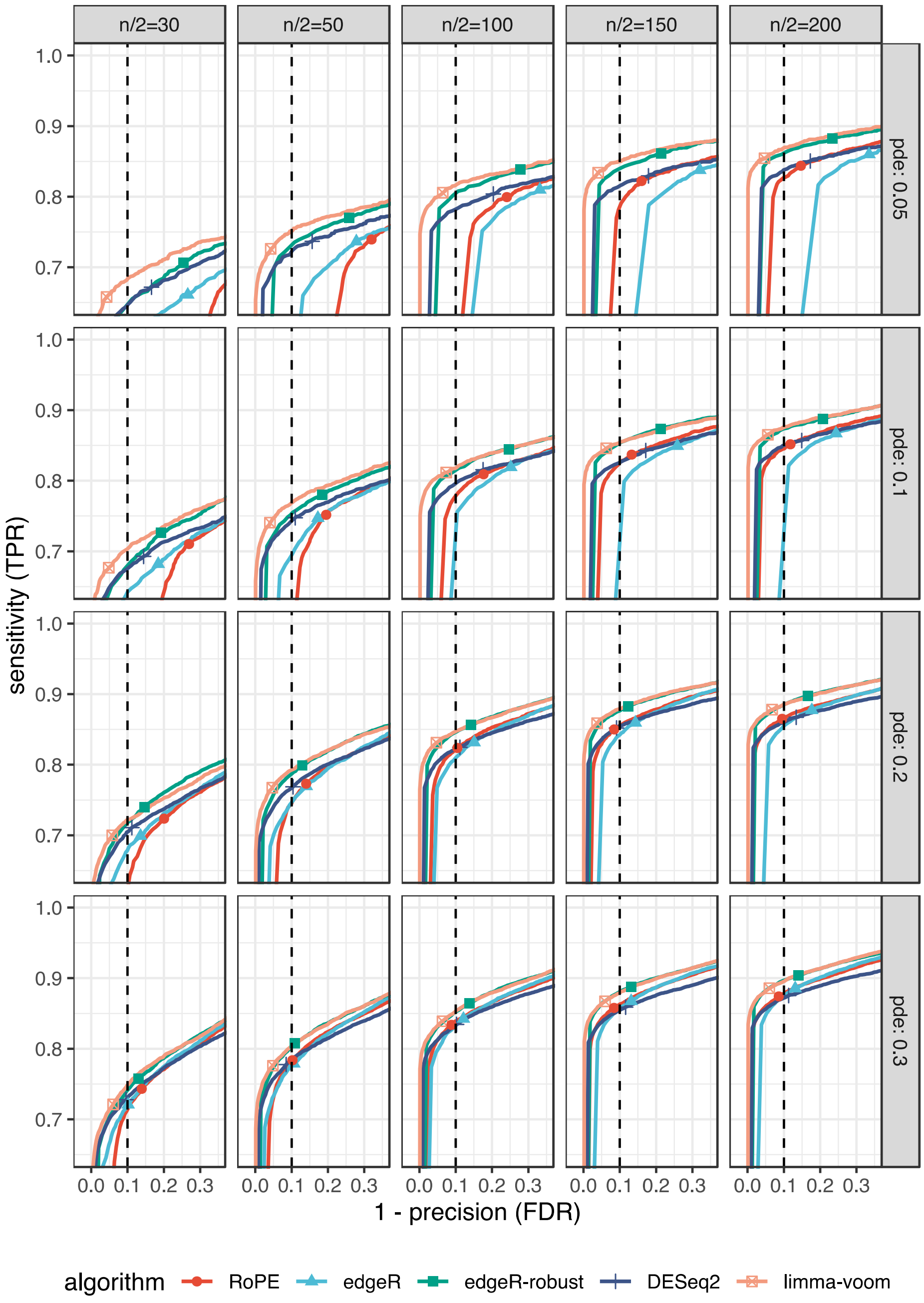


**Figure S6: Non-parametric simulation result varying the percentage of true DE gene from 5% to 30%: TPR vs FDR**

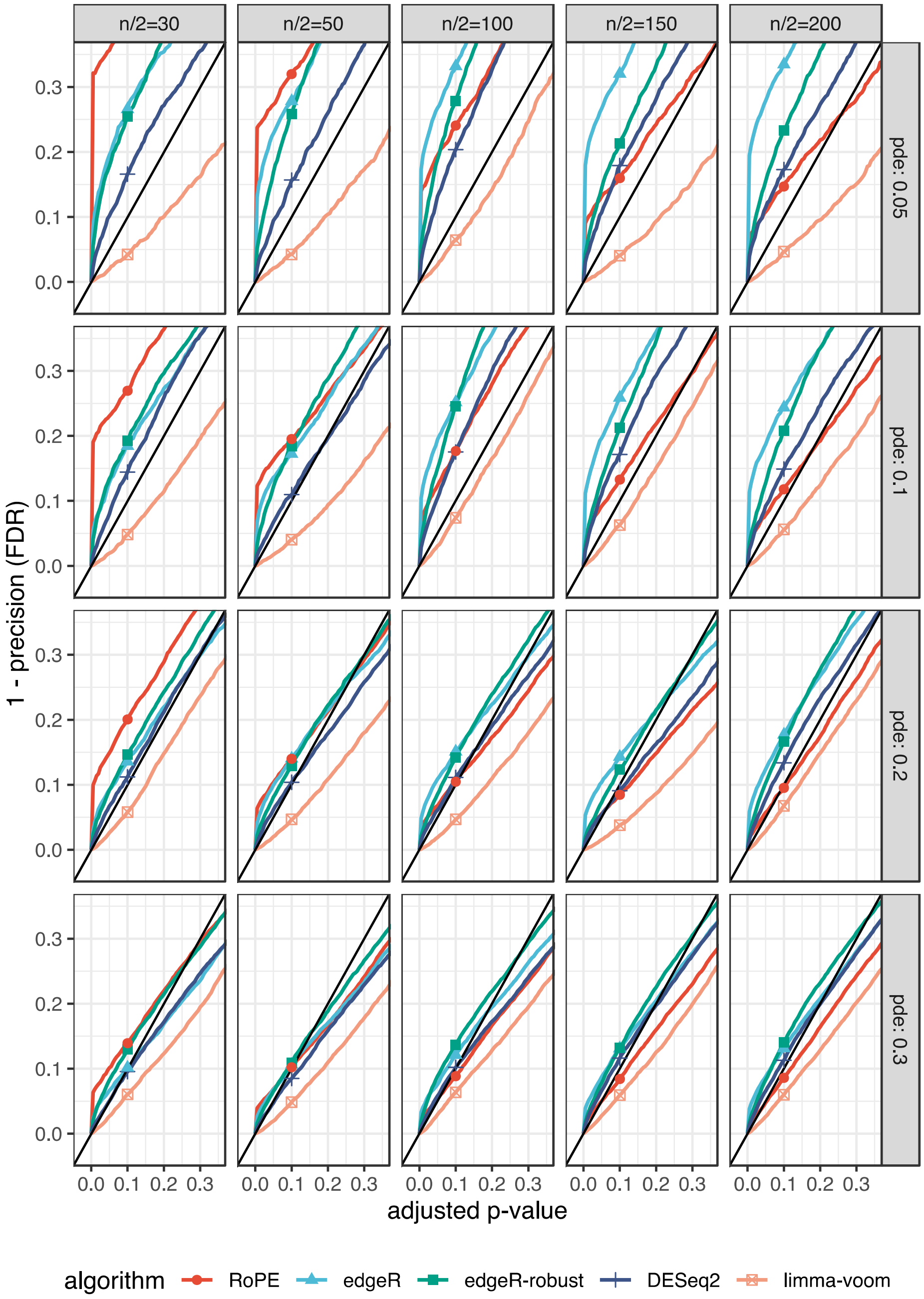


**Figure S7: Non-parametric simulation result varying the percentage of true DE gene from 5% to 30%: FDR vs Adj-Pvalue**

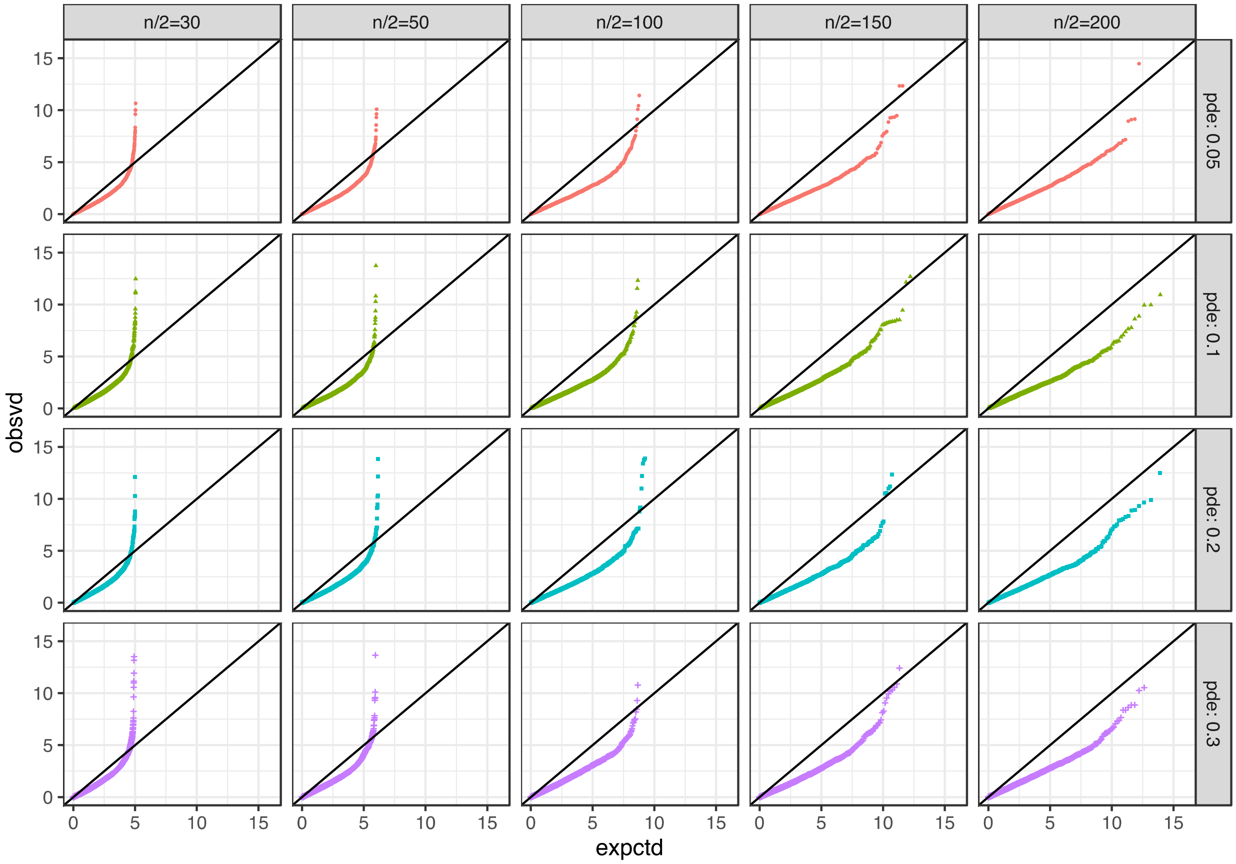


**Figure S8: Chi squared (with df = 1) quantile-quantile plot of RoPE test statistics under parametric simulation.**

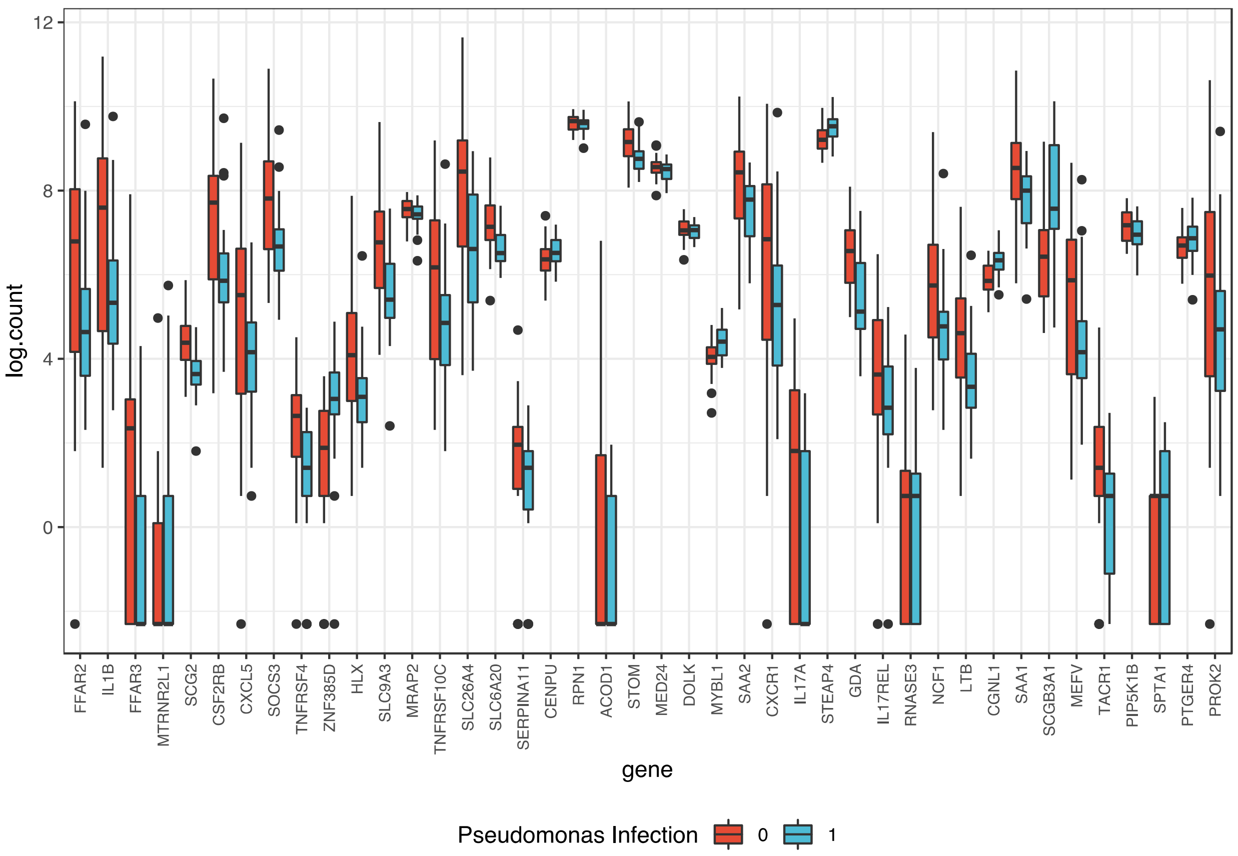


**Figure S9: Boxplot of protein-coding DE gene expression ranked by p-value (smallest on the left) for CF patients with vs without PsA infection.** The expression is shown as the natural logarithm of (RNA-Seq count + 0.01). All genes are identified as differentially expressed by the RoPE tool (adjusted p-value < 0.05). SLC9A3 and SLC26A4 were ranked at 16 and 20.

| Method | logFC | p-value | Adj p-value | Rank |
|---|---|---|---|---|
| *edgeR* | -1.5348 | 0.001300 | 0.929677 | 41 |
| *DESeq2* | -1.0604 | 0.000926 | 0.678998 | 40 |
| *voom* | -1.7561 | 0.004593 | 0.999784 | 84 |
| *RoPE* | -1.3917 | 0.000003 | 0.004204 | 20 |

**Table S1: Summary of differential expression analysis of SLC26A4 gene.** For each method, this summary table contains the estimates of natural logarithm of differential expression fold changes (logFC), the nominal p-values, the BH adjusted p-values (Adj p-value) for DE tests, and the rank of DE significance by adjusted p-values for SLC26A4.

#### Table S2 Top DE genes with adjusted p-values less than 0.05, ranked by RoPE

| Symbol | logFC | Adjustment Factor | p-value | Adj p-value | edgeR Rank | DESeq2 Rank | voom Rank |
|---|---|---|---|---|---|---|---|
| FFAR2 | -0.9243 | 0.0055 | <0.000001 | <0.000001 | 710 | 775 | 80 |
| IL1B | -0.9072 | 0.0023 | <0.000001 | <0.000001 | 537 | 510 | 541 |
| FFAR3 | -0.6901 | 1.1663 | <0.000001 | <0.000001 | 36 | 210 | 5,349 |
| MTRNR2L1 | 3.6284 | 0.0690 | <0.000001 | 0.000002 | 9,681 | 10,812 | 3,788 |
| AC098476.1 | -0.7887 | 0.1114 | <0.000001 | 0.000051 | 4 | 3 | 2 |
| RPL10P6 | 2.3654 | 0.0700 | <0.000001 | 0.000051 | 160 | 113 | 4,035 |
| SCG2 | -0.9086 | 0.0444 | <0.000001 | 0.000083 | 1 | 1 | 1 |
| CSF2RB | -0.4180 | 0.0063 | <0.000001 | 0.000468 | 349 | 433 | 21 |
| CXCL5 | -1.3391 | 0.0038 | <0.000001 | 0.000468 | 89 | 92 | 494 |
| SOCS3 | -0.6369 | 0.0019 | 0.000001 | 0.002167 | 68 | 83 | 108 |
| AC006480.1 | 1.0168 | 1.8335 | 0.000001 | 0.003517 | 107 | 511 | 1,814 |
| AC022826.1 | 1.4359 | 0.5789 | 0.000001 | 0.003590 | 9 | 16 | 7 |
| TNFRSF4 | -0.7277 | 0.3606 | 0.000002 | 0.004141 | 21 | 26 | 145 |
| ZNF385D | 1.0423 | 0.0924 | 0.000002 | 0.004141 | 14 | 17 | 12 |
| HLX | -0.6342 | 0.0528 | 0.000002 | 0.004141 | 2,947 | 3,488 | 1,732 |
| SLC9A3 | -1.2705 | 0.0011 | 0.000002 | 0.004141 | 8 | 9 | 92 |
| MRAP2 | -0.2094 | 0.0205 | 0.000003 | 0.004141 | 17 | 11 | 5 |
| AL138789.1 | 1.1594 | 0.1991 | 0.000003 | 0.004141 | 680 | 967 | 211 |
| TNFRSF10C | -0.4534 | 0.0134 | 0.000003 | 0.004141 | 2,169 | 2,709 | 3,213 |
| SLC26A4 | -1.3917 | 0.0002 | 0.000003 | 0.004204 | 41 | 40 | 84 |
| SLC6A20 | -0.5985 | 0.0039 | 0.000004 | 0.005561 | 6 | 8 | 10 |
| SERPINA11 | -1.1857 | 0.1566 | 0.000005 | 0.005850 | 25 | 27 | 1,075 |
| CENPU | 0.3430 | 0.0199 | 0.000005 | 0.005850 | 15 | 12 | 4 |
| MTND1P23 | -2.9151 | 0.0012 | 0.000005 | 0.006198 | 339 | 75 | 10,505 |
| RPN1 | -0.1001 | 0.0104 | 0.000006 | 0.006757 | 271 | 18 | 15 |
| AC022509.2 | 1.2849 | 0.5397 | 0.000006 | 0.007218 | 38 | 67 | 18 |
| RNU2-33P | -1.0409 | 1.1687 | 0.000007 | 0.008098 | 40 | 80 | 220 |
| ACOD1 | -2.3659 | 0.0584 | 0.000010 | 0.010441 | 17,222 | 11,225 | 20,858 |
| STOM | -0.3988 | 0.0011 | 0.000011 | 0.011122 | 10 | 13 | 11 |
| MED24 | -0.1506 | 0.0127 | 0.000011 | 0.011232 | 44 | 30 | 16 |
| DOLK | -0.1201 | 0.0846 | 0.000012 | 0.011345 | 116 | 28 | 25 |
| MYBL1 | 0.3802 | 0.1372 | 0.000014 | 0.012465 | 20 | 7 | 8 |
| AL079342.3 | 1.0419 | 0.4250 | 0.000017 | 0.014831 | 28 | 37 | 29 |

| | | | | | | | |
|---|---|---|---|---|---|---|---|
| SAA2 | -0.8153 | 0.0006 | 0.000018 | 0.015406 | 80 | 64 | 103 |
| CXCR1 | -0.5843 | 0.0026 | 0.000021 | 0.017827 | 2,705 | 2,951 | 6,969 |
| IL17A | -1.3840 | 0.0814 | 0.000022 | 0.017999 | 789 | 701 | 143 |
| STEAP4 | 0.3131 | 0.0011 | 0.000023 | 0.018586 | 22 | 20 | 6 |
| AL355499.1 | -0.9780 | 0.8121 | 0.000030 | 0.023135 | 247 | 262 | 422 |
| GDA | -0.7861 | 0.0038 | 0.000032 | 0.023811 | 2 | 2 | 3 |
| PICART1 | 0.3797 | 0.2851 | 0.000034 | 0.024854 | 78 | 68 | 48 |
| IL17REL | -1.1193 | 0.0184 | 0.000036 | 0.025085 | 7 | 6 | 1,074 |
| SLC26A4-AS1 | -0.7463 | 0.0024 | 0.000036 | 0.025085 | 52 | 74 | 134 |
| RNASE3 | 1.2288 | 0.2944 | 0.000037 | 0.025085 | 1,547 | 2,300 | 1,989 |
| RNU4ATAC18P | 0.8950 | 2.2011 | 0.000048 | 0.031777 | 194 | 1,708 | 94 |
| NCF1 | -0.3733 | 0.0208 | 0.000049 | 0.031777 | 1,302 | 1,819 | 2,756 |
| LTB | -0.3115 | 0.1246 | 0.000054 | 0.034594 | 2,400 | 2,739 | 2,642 |
| CGNL1 | 0.3200 | 0.0248 | 0.000055 | 0.034594 | 27 | 24 | 13 |
| SAA1 | -0.8713 | 0.0003 | 0.000057 | 0.035108 | 152 | 152 | 299 |
| SCGB3A1 | 1.0269 | 0.0004 | 0.000060 | 0.035252 | 11 | 14 | 9 |
| MEFV | -0.5107 | 0.0107 | 0.000061 | 0.035252 | 768 | 957 | 1,469 |
| TACR1 | -1.4288 | 0.0924 | 0.000061 | 0.035252 | 43 | 45 | 409 |
| GRIK1-AS1 | 1.0314 | 0.1005 | 0.000068 | 0.038078 | 5 | 5 | 32 |
| PIP5K1B | -0.2574 | 0.0145 | 0.000071 | 0.039033 | 29 | 46 | 20 |
| FAM133CP | -1.0037 | 0.2339 | 0.000073 | 0.039839 | 313 | 240 | 414 |
| SNORD89 | -0.5646 | 1.0016 | 0.000081 | 0.043359 | 828 | 1,998 | 1,596 |
| AC034105.3 | 1.1360 | 0.8418 | 0.000083 | 0.043567 | 254 | 1,505 | 1,578 |
| SPTA1 | 0.9012 | 0.5465 | 0.000085 | 0.043885 | 707 | 865 | 1,284 |
| PTGER4 | 0.3367 | 0.0111 | 0.000092 | 0.046538 | 45 | 54 | 49 |
| PROK2 | -0.6360 | 0.0035 | 0.000097 | 0.048108 | 29,103 | 27,928 | 10,462 |
